# Estimating the correlation of exchangeable variables in assortative mating

**DOI:** 10.64898/2026.08.22.746446

**Authors:** Gabriel Kennedy, Alejandro Ochoa

**Affiliations:** Duke University Program in Genetics and Genomics, Duke University, Durham NC 27705, USA; Department of Human Genetics, University of California, Los Angeles, Los Angeles, CA 90095, USA; Department of Biostatistics and Bioinformatics, Duke University, Durham, NC 27705, USA; Duke Center for Statistical Genetics and Genomics, Duke University, Durham, NC 27705, USA

**Keywords:** assortative mating, admixture, exchangeable variables, Pearson, correlation, order bias

## Abstract

In studies of assortative mating, similarity between variables measured in parents is often quantified using correlation. The order of the parents within any given pair can be arbitrary in these applications, but common correlation estimators are not robust to reordering within pairs. These unordered variable pairs are exchangeable, since the joint distributions of both orders are equal, and a given order is biased if the one variable has a lower expectation than the other. In this work, we characterize the effect of order bias on Pearson correlation estimates assuming exchangeable variables, and develop a new unbiased estimator, CorSym, that does not depend on order within each pair. Exchangeable variables have equal marginal distributions for both variables, a property accounted for by CorSym. In contrast, standard correlation estimators assume the two variables have different distributions, so biased orders skew the underlying mean, variance and covariance estimates. We show, through theory and simulations, how order bias often results in upwardly biased Pearson correlation estimates. Simulations confirm CorSym is unbiased, and validate its estimated confidence intervals. Using real admixed trios (parents and a child) from 1000 Genomes, we first demonstrate that the global ancestry of fathers and mothers are consistent with exchangeability, using both Kolmogorov-Smirnov tests and a Binomial test for order bias. However, ANCESTOR, which estimates parental global ancestry from a child’s local ancestry, produces significant order biases in its output that result in substantial Pearson biases, which CorSym overcomes. Compared to ancestry proportions calculated directly on the parents, ANCESTOR also overestimates parent ancestry divergence and experiences another estimation artifact. Overall, CorSym solves an important estimation bias likely to be encountered in the study of assortative mating, providing unbiased and deterministic estimates that do not depend on the arbitrary order of the data.

## 1 Introduction

Ordinary correlation estimation between two variables assumes that they are different in essence, an asymmetry that manifests in differences in their marginal means and variances. In this general case you cannot swap values in a given pair but not in other pairs, since they could have different units (for example, income and age), but even variables with the same units can have different distributions (such as height and waist circumference). Here, we consider the correlation of two measurements of the same variable where the order of the measurements is arbitrary; for example, income in a couple. In statistics, such variable pairs are called exchangeable. As we show in this work, the added flexibility of ordering within each pair causes new problems not observed for non-exchangeable variables. In particular, Pearson estimates differ depending on this order, and upwardly biased estimates arise often if the order is biased: if the first value is more likely to be smaller than the second one, or vice versa.

Assortative pairing is a key application where correlation is estimated between pairs of exchangeable variables. For example, consider whether individuals with more similar incomes are more likely to be paired, where pairing can refer to dating, marriage, cohabitation, or sharing children. For opposite sex pairs of known sex, sex could order the pairs (for example, values for women first), but this is not possible when including same-sex pairs or when sex is unknown. Individuals could be ordered by other variables, such as age (for example, oldest first), though as we show, different orders change Pearson estimates, so any such ordering must be justified. In the worst case, in seeking to order data uniquely without separate ordering variables, one might always list the lower value first (or last), which we show here maximizes correlation bias.

Assortative mating in humans is a type of non-random mating favoring similar sociodemographics, age, health, education, language, and other variables [1, 2, 3, 4, 5]. In ancestry-based assortative mating, mate choice depends on global (genome-wide average) genetic ancestry similarity. In recently admixed populations such as Hispanics and African-Americans—whose individuals have African, European and Native American ancestries in variable proportions—there are positive correlations between the major ancestries of pairs [6, 7, 8, 9]. However, sample information for ordering parents is often unavailable due to limited data collection or privacy protections. Recent methods, including ANCESTOR, attempt to infer parental global ancestry using only their child’s local ancestry (predicted over genomic regions) [10, 11, 12, 13]. These methods output parents in an arbitrary (and in at least one case, a biased) order, without additional information to order parents meaningfully.

In this work, we develop a new correlation estimator for exchangeable variables, constructed to be invariant to the order within each pair. We call this estimator “CorSym” since it assumes a symmetry between the two variables. We characterize the bias in Pearson estimation due to order bias and validate CorSym using theory and simulations. We then employ trio samples from the latest 1000 Genomes Project [14] to validate our exchangeability assumption and demonstrate that the parent ancestries output by the inference method ANCESTOR have an order bias that carries over to correlation bias with Pearson but not CorSym. We recommend CorSym for estimating correlations between exchangeable variables, which arise in assortative pairing, to correctly model arbitrary order and ensure reproducible, deterministic estimates.

## 2 Methods

### 2.1 Definitions

The variables *X* and *Y* are exchangeable if the distribution of (*Y, X*) is identical to that of (*X, Y*), so their marginal distributions are identical:

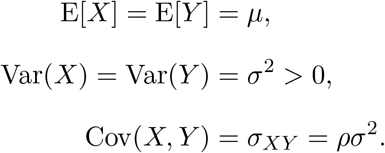

We will treat different pairs as independent.

More generally, the pair (*X, Y*) is said to have a “random” order if E[*X*] = E[*Y*], and its order is “biased” if E[*X*]≠ E[*Y*]. Pairs with a biased order become “randomized” if individual pairs (*X, Y*) are randomly replaced by (*Y, X*) independently with probability 1/2. Lastly, the order is “extreme” if *X* ≤ *Y* always holds (or *X* ≥ *Y* always). Note that extreme order is a special case of biased order. In this work, simulated data is always drawn in random order, and we impose biased orders afterwards, whereas real data starts out in a possibly biased order, and we subsequently randomize it for our comparisons.

In simulated data with random order, the forced ordering proportion *q* is defined as the proportion of pairs (*X, Y*) replaced with their extreme order counterparts (*L, H*), where

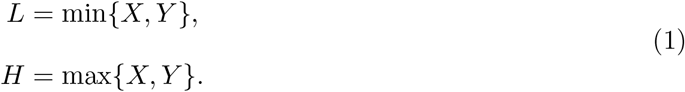

The resulting order proportion, which is the expected proportion of times that *X* ≤ *Y*, is *f* = 0.5(1 − *q*) + *q* = 0.5 + *q/*2, and it is said to be biased if *f*≠ 0.5.

### 2.2 Theoretical correlation upward bias under extreme ordering

Here we characterize an extreme case, with the understanding that partial orderings will exhibit intermediate biases. Compare the distribution of (*X, Y*) to the extreme ordering pair (*L, H*) defined earlier. Eq. (1) suggests that *L* and *H* have an additional dependence on each other compared to the original pair, since both depend on both of the input variables *X* and *Y*. Letting E[*L*] = *µ*_*L*_, Var(*L*) = *σ*^2^, Var(*H*) = *σ*^2^, and Cor(*L, H*) = *ρ*_*LH*_, in Appendix A, we prove that

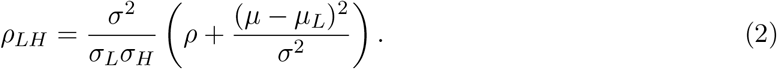

Furthermore, *ρ*_*LH*_ ≥ *ρ* holds if *ρ*_*LH*_ ≥ 0, and also if *σ*_*L*_ = *σ*_*H*_ (guaranteed if *X* has a symmetric distribution). Thus, the correlation of extreme ordered variables is greater than that of the original exchangeable variables in most cases. Conversely, *ρ*_*LH*_ ≤ *ρ* requires both negative *ρ*_*LH*_ and a skewed marginal distribution for *X*. Pearson estimates of extremely ordered data will approach *ρ*_*LH*_ for large sample sizes.

For example, consider *X, Y* ~ Uniform(0, 1) drawn independently. Their moments are well known to be *µ* = 1*/*2, *σ*^2^ = 1*/*12, and *ρ* = 0. Furthermore, their minimum *L* and maximum *H* are the well-character order statistics of the uniform distribution, which have Beta distributions. Their marginal moments are *µ*_*L*_ = 1*/*3, *µ*_*H*_ = 2*/*3, and 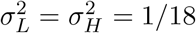. Applying these values to Eq. (2) we obtain *ρ*_*LH*_ = 1*/*2, which is remarkably large even though we started from independent variables.

The positive correlation between *L* and *H* makes sense, since when *H* exceeds its expectation, that gives *L* more room above its own expectation. Conversely, when *H* is lower than its expectation, it restricts *L* more (since *L* ≤ *H*), making it more likely to be smaller than its expectation. Thus, the residuals of *L* and *H* from their expectations are more likely to have the same sign, so they are more correlated, compared to the starting variables *X* and *Y*.

### 2.3 Correlation estimator for exchangeable variables: CorSym

We use the method of moments to derive the new correlation estimator for exchangeable variables, which by construction gives the same answer when elements in any pair are swapped. We first define “base” variance and covariance estimators, which are biased for small samples, then construct unbiased estimators.

Let the observed *N* pairs be 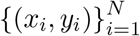. The mean 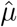 and base variance 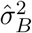 estimators below are standard sample estimates applied to the pooled data (combining both values of each pair, totaling 2*N* samples). The base covariance estimator 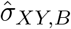 is the sample estimator but using the pooled mean estimate, which forces a different normalization factor 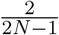 instead of the usual Bessel’s correction of 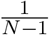):

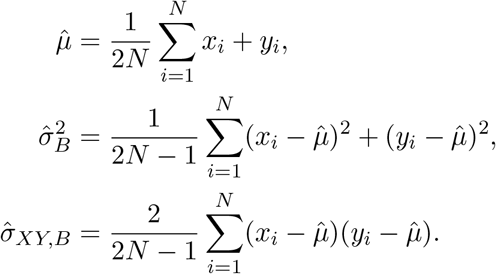

Although 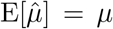 holds, the base variance and covariance estimators biased for small samples (calculations in Appendix B):

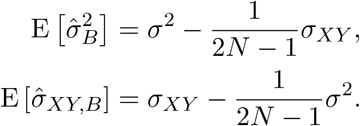

We solve this system of equations to obtain unbiased estimates (derivation in Appendix C):

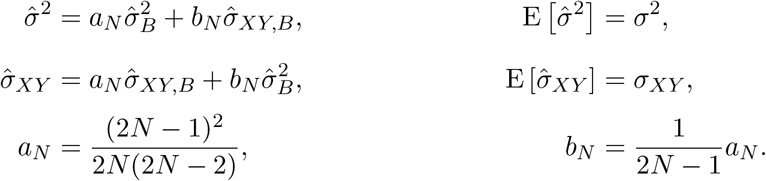

Note that for large samples *a*_*N*_ ≈ 1 whereas *b*_*N*_ vanishes. Lastly, the correlation is estimated by the ratio of the unbiased covariance and variance, which estimates the true correlation with a small bias that ratio estimators have:

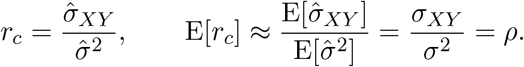

Note that every component of this estimator is invariant to swapping *x*_*i*_ and *y*_*i*_ for the same *i*, for any subset of such pairs *i*, as desired.

### 2.4 Pearson correlation estimator

The Pearson estimator is given by

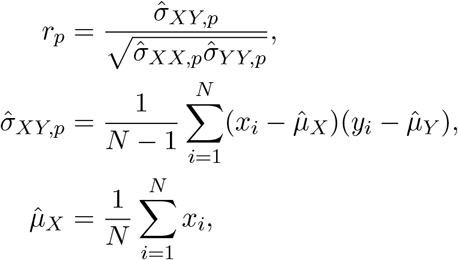

where 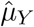 is like 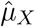 but replacing *x*_*i*_ with *y*_*i*_, and so on. Assuming the variables are exchangeable and in random order, the mean, variance and covariance estimates are unbiased because their assumptions are met, so this estimator is also approximately unbiased. However, this approach gives different estimates if *x*_*i*_ and *y*_*i*_ are swapped in a given pair *i*. A biased order leads to biases in every component of this estimator (means, variances, and covariance) and the final correlation estimate, as illustrated earlier for extreme ordering.

### 2.5 Confidence intervals (CIs)

CorSym estimates CIs using the same strategy as Pearson. First, the estimated correlation *r* undergoes the Fisher transformation, which is the inverse hyperbolic tangent [15]:

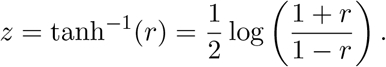

When *r* is a Pearson estimate and the input *X* and *Y* are Multivariate Normal (and for large samples due to the Central Limit Theorem), this *z* has an approximate Normal distribution with mean equal to the transformed *ρ* and a variance of 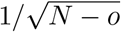, where the offset is *o* = 3 for Pearson. For CorSym this approach also works except *o* = 1.5 performs best (see Results). Based on that theory, CIs at the *α* level (which cover the true parameter with probability 1 − *α*) are estimated by

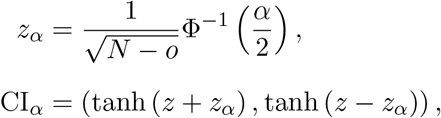

where Φ is the Standard Normal cumulative function.

### 2.6 Order bias test

Let *c* be the number of pairs with *X* ≤ *Y* (see treatment of ties shortly), and 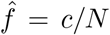 be the estimated order proportion. Under the null hypothesis of no order bias, the counts have a Binomial distribution with probability of success *f*_0_ = 0.5:

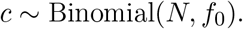

Two-sided p-values are calculated under this distribution.

In practice, ties were observed frequently, especially for minor ancestries (when both equal zero or some other small value, due to limited precision). Counting as either *X* ≤ *Y* or *X < Y* assigns all ties to a single order, incorrectly causing an order bias. Therefore, we count ties as half observations for both orders: they each contribute 1 to *N* and 0.5 to *c*, the final *c* is rounded down if not integer. Thus, ties are treated as observations favoring the null hypothesis.

### 2.7 Simulations for estimator evaluation

100 replicates with bivariate Normal distribution were simulated using *mvrnorm* from the R package *MASS*, all with zero mean and unit variance, for each *ρ* from −1 to 1 by 0.01 and sample sizes *N* from 10, 50, 100, 500, and 1000 [16].

Non-normal data were simulated using the *mvdc* function from the *copula* R package [17, 18, 19, 20]. A copula produces correlated Uniform variables, which can be transformed into any desired marginal distribution. Data were simulated for *ρ* in −1, −.5, 0, .5, and 1, with equal Beta marginal distributions for exchangeability. Tested Beta distributions included uniform (*α* = *β* = 1), skewed (*α* = 1, *β* = 8), and U-shaped (*α* = *β* = 0.5).

Each pair has two orders, the original random order and the extreme order where the higher value in each pair is always listed first. In our forced ordering proportion analysis, proportions *q* of 0%, 25%, 50%, 75%, and 100% were considered, where a random proportion *q* of pairs is replaced with their extreme order (0% is random order and 100% is extreme order).

### 2.8 Empirical Data Analysis

We used 170 unique trios from 6 admixed populations (PEL, *N* = 35 trios; PUR, *N* = 35; MXL, *N* = 32; CLM, *N* = 35; ASW, *N* = 13; ACB, *N* = 20) from the 1000 Genomes Project [14]. For ancestry references, we used non-trio samples from the merged Human Genome Diversity Project (HGDP) and 1000 Genomes [21, 22, 23, 24]. In particular, for African and European ancestry we used the YRI (*N* = 175) and IBS (*N* = 151) populations from 1000 Genomes, respectively, whereas for Native American ancestry (labeled NAM, *N* = 62) we merged the Pima, Maya, Colombian, Surui, and Karitiana populations from HGDP. Three-way ancestry analyses included IBS-like, YRI-like, and NAM-like ancestries, whereas 2-way analyses compared YRI-like to Non-YRI-like (combined IBS-like and NAM-like). We retained only autosomal SNPs, and filtered trios and references separately to keep SNPs with a minor allele frequency greater than 0.01 and missingness less than 0.1, then trios and references were merged at the intersection of loci, using `plink2` [25]. Trios and references were phased separately using BEAGLE 5.5 [26].

Gold standard ancestry proportions were estimated using the ADMIXTURE software [27] applied to the merged reference populations and trios (children and parents). We treat these estimates as true global ancestries, since their accuracy is very high. Prior to admixture analysis only, SNPs were LD pruned with `plink2 --indep-pairwise` to ensure all SNP pairs over a 50kb window satisfy 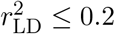. Kolmogorov-Smirnov tests were used to compare mother and father ancestry proportion distributions, to test our exchangeability assumption.

For parental ancestry inference, local ancestry blocks were inferred in trio children only using RFMIX [28]. Lastly, parental global ancestries were inferred from child local ancestry (ignoring the parent samples) using ANCESTOR [10].

## 3 Results

### 3.1 Simulation evaluations

We simulate exchangable pairs with known correlations to characterize the bias in the Pearson estimator that CorSym solves. First we simulated *N* = 1, 000 Standard Normal variable pairs with true correlations of −1, −.5, 0, .5, and 1, and calculated both *r*_*c*_ and *r*_*p*_. With data in random order, *r*_*c*_ and *r*_*p*_ agree (first row of Fig. 1A). However, with extreme order (the smallest of the two values listed first) *r*_*p*_ incorrectly overestimates the correlation, while *r*_*c*_ is unaffected and thus remains unbiased (second row of Fig. 1A). As correlations approach 1 or −1, bias is reduced to zero. In non-Normal data generated using copulas with skewed, Uniform, and U-shaped marginal distributions, *r*_*p*_ experiences strong biases under extreme order, whereas *r*_*c*_ is unaffected (Fig. S1).

**Figure 1.**
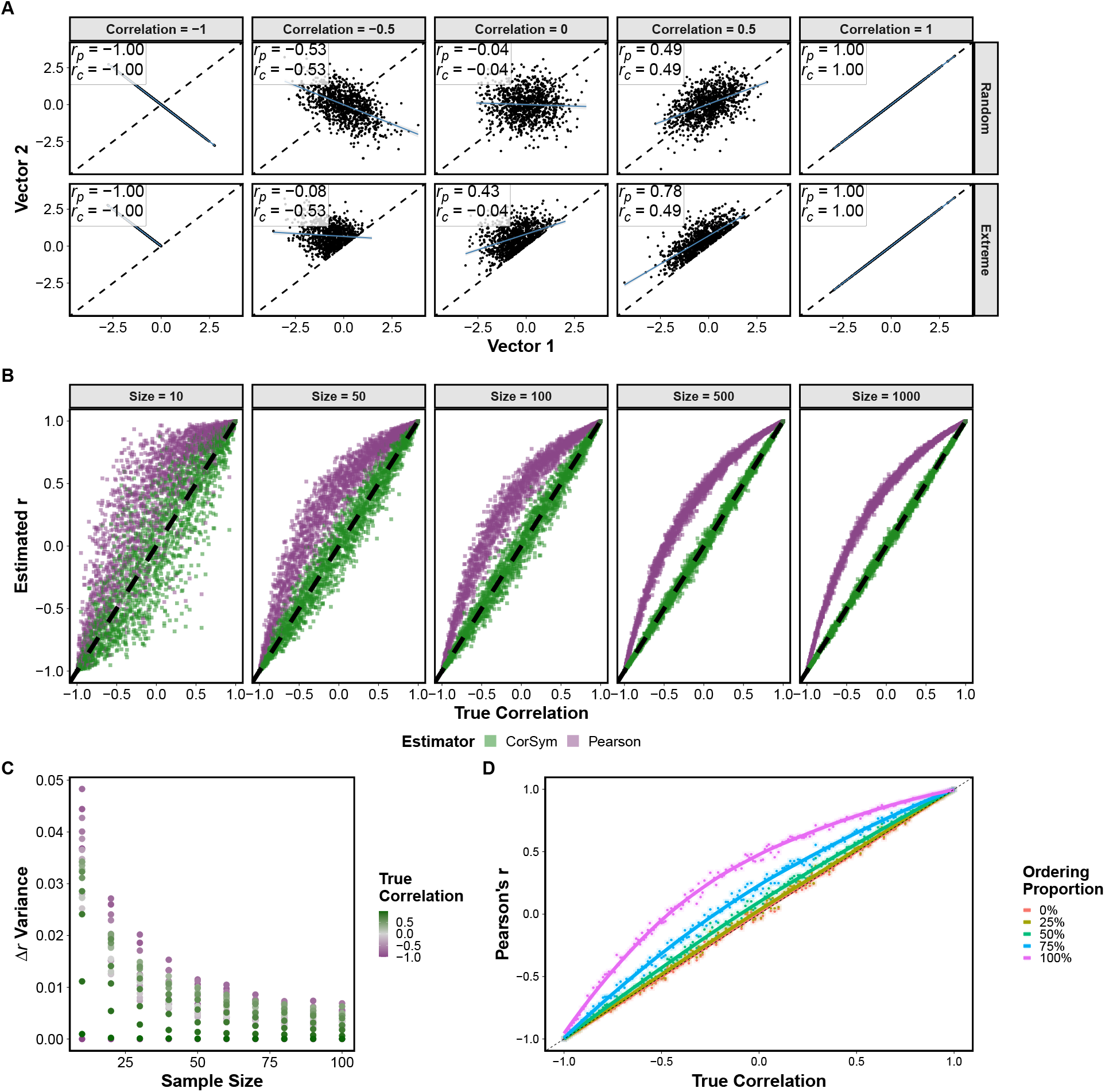
Testing CorSym and Pearson with simulated correlated Normal pairs. **A)** Exchangeable data was simulated (*N* = 1000) for several *ρ* values (top row). Extreme order forces all values of vector 1 to be lower than those of vector 2 (bottom row). **B)** CorSym and Pearson estimates of extreme ordered data computed for 100 replicates per *ρ* between −1 and 1 by .01 increments, for 5 sample sizes. **C)** The variance of the difference between CorSym and Pearson estimates (Δr) decreases with sample size. **D)** Pearson’s bias increases with the forced ordering proportion *q* (*N* = 1000).

We then simulated 100 replicates with extreme order for each true *ρ* (−1 to 1, by 0.01) and sample size combination (10, 50, 100, 500, 1,000). At 10 samples, despite noise in the data and high variance, there is a noticeable bias in *r*_*p*_, while *r*_*c*_ is unbiased (Fig. 1B). As sample sizes increase, estimation variance decreases, revealing the functional form of Pearson’s upward bias, which is greatest near *ρ* = 0. The variance of *r*_*c*_ − *r*_*p*_ decreased with sample size and as *ρ* approached −1 or 1 (Fig. 1C). Since the previous simulations have no marginal skew, they result in upward biases only, as theory predicted. We also simulated highly skewed log-Normal data to demonstrate downward biases for very negative correlations, all together expected to be uncommon in practice (Fig. S2).

Next, we considered partial orderings. Forced ordering proportions *q* of 0% (random order), 25%, 50%, 75% and 100% (extreme) were compared at *N* = 1, 000. As expected, the bias of Pearson estimates increases with the forced ordering proportion (Fig. 1E). Interestingly, bias itself increases non-linearly with *q*, with the greatest increase seen between 75% and 100% ordering proportions, whereas bias is marginal at 25% ordering compared to 0% ordering.

Lastly, we validate CorSym’s 95% confidence intervals (CIs), which are calculated with a formula derived for Pearson. Simulations for a given *N* and *ρ* were replicated one million times to calculate CI coverage (proportion of replicates where CIs contained the true correlation parameter, ideally 0.95). First, using *ρ* = 0 and *N* = 10, we tested offset parameters *o* between 0 and 3 (Pearson uses *o* = 3), and *o* = 1.5 gave the most calibrated CIs for CorSym (Table S1). Using *o* = 1.5 from now on, we confirmed calibrated CI coverages for all sample sizes and true correlations tested (Fig. S3). Thus, CorSym CIs are well calibrated, even at the smallest sample sizes considered.

### 3.2 Empirical Evaluations

Here we show that the inference software ANCESTOR outputs parental ancestries with a biased order, and demonstrate the resulting biased correlations estimates using Pearson, which CorSym overcomes. We use admixed trios from 1000 Genomes to demonstrate this behavior, which also allows us to compare inferred to true parent ancestries (estimated directly from parent genomes using the software ADMIXTURE, instead of from the child only as ANCESTOR does). This allows us to characterize additional inference errors unrelated to order bias but equally consequential to correlation estimation.

#### 3.2.1 Father and mother true ancestry distributions are exchangeable

First we establish that parent true ancestry proportions (high accuracy ADMIXTURE estimates) are exchangeable by comparing the distributions of mothers and fathers. We tested *K* = 2 to 6 ancestries using cross validation and confirmed the best fit at *K* = 3, matching the number of reference populations included in this analysis (Fig. S4). The ancestry distribution of the admixed trios and reference panels is shown in Fig. 2A. We found no significant differences between the ancestry distribution of mothers and fathers of each population at any of the three ancestries (Fig. 2B; Kolmogorov-Smirnov p-values in Table S2). We also performed biased order tests for each population and ancestry, as well as a test combining populations, none of which were significant (Table 1). We caution that these populations have small sample sizes, so we may not have enough power to detect differences. Nevertheless, we found no evidence for parents ordered by sex having different true ancestry distributions, so treating them as exchangeable is adequate.

**Table 1.** True ancestry order proportions ordered by sex.

| Population | $N$ | YRI-like | | | IBS-like | | | NAM-like | | |
| --- | --- | --- | --- | --- | --- | --- | --- | --- | --- | --- |
| | | $F \geq M$ | $\hat{f}$ | P-value | $F \geq M$ | $\hat{f}$ | P-value | $F \geq M$ | $\hat{f}$ | P-value |
| ACB | 20 | 12 | 0.6 | 0.503 | 8 | 0.4 | 0.503 | 9 | 0.45 | 0.824 |
| ASW | 13 | 6 | 0.462 | 1 | 8 | 0.615 | 0.581 | 5 | 0.385 | 0.581 |
| CLM | 35 | 15 | 0.429 | 0.5 | 19 | 0.543 | 0.736 | 19 | 0.543 | 0.736 |
| MXL | 32 | 13 | 0.406 | 0.377 | 15 | 0.469 | 0.86 | 15 | 0.469 | 0.86 |
| PEL | 35 | 16 | 0.457 | 0.736 | 17 | 0.486 | 1 | 18 | 0.514 | 1 |
| PUR | 35 | 15 | 0.429 | 0.5 | 20 | 0.571 | 0.5 | 18 | 0.514 | 1 |
| Total | 170 | 77 | 0.453 | 0.25 | 87 | 0.512 | 0.818 | 84 | 0.494 | 0.939 |

F=Father, M=Mother. No tests were significant at 0.05 level after Bonferroni correction for 21 tests ( $p < 0.00238$ ).

**Figure 2.**
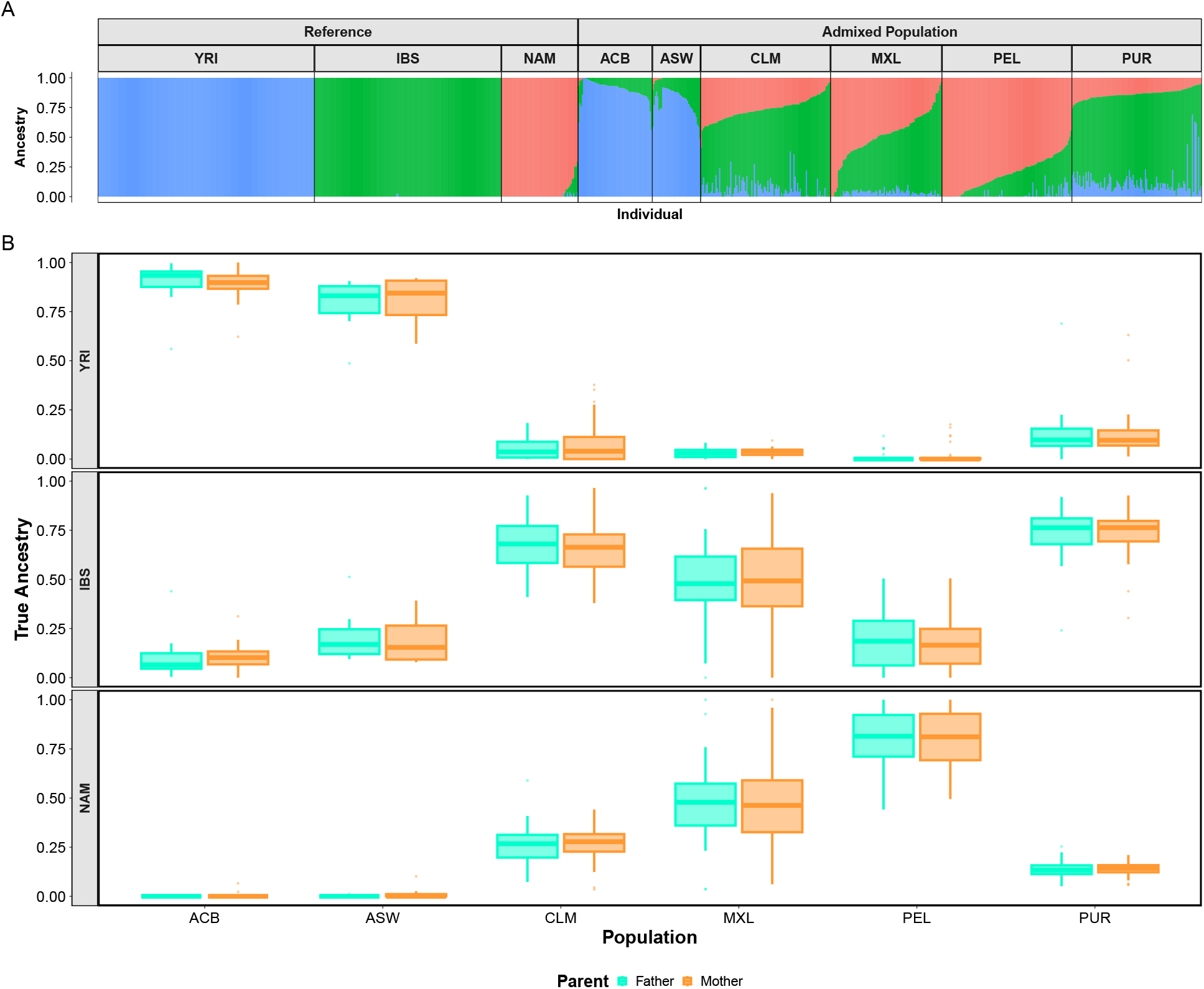
Mother and father ancestry distributions are consistent with exchangeability. Ancestry proportions (y-axis) estimated on all samples directly using ADMIXTURE are treated as true ancestry. **A)** Ancestry proportions of reference individuals and admixed trios (children and all parents; x-axis). **B)** Parent ancestry proportion distributions were not significantly different between sexes by Kolmogorov-Smirnov test (p-values in Table S2) for YRI-like, IBS-like, and NAM-like ancestries.

#### 3.2.2 ANCESTOR outputs have biased order

In contrast to true parent ancestries ordered by sex, here we show that outputs from the inference software ANCESTOR have a biased order. ANCESTOR does not predict the sex of the parents or any other separate ordering variables, so the order of the parents (P1, P2) is presumably arbitrary. However, in two-way admixture inference (YRI-like vs Non-YRI-like ancestries), ANCESTOR tended to assign P1 higher YRI-like ancestry than P2 (Fig. 3A). All ASW parent pairs are ordered extremely, followed by ACB with 80% biased order proportion, while the smallest was for PEL with 60% (Table 2). These proportions are usually not significantly different from random order due to small sample sizes, except in ASW, but the bias is significant after pooling populations: 69% of the time the first parent has greater YRI-like ancestry (two-sided Binomial *p* = 1.02e-6). We next randomized 100 times the parental order of the ANCESTOR estimates. As desired, randomly ordered samples were better balanced on which parent was assigned the higher ancestry (Fig. 3B). True ancestries ordered by sex are included for comparison in Fig. 3C. CorSym estimates on the original and randomized orders were numerically identical, also as expected (Fig. 3D). CorSym estimates are also similar to the Pearson estimates in each randomization and they differ greatly from Pearson estimates in the original order, a bias correlated to the ordering proportion, except in MXL, a population with limited YRI ancestry (Fig. 3E, Table 2). Two cases have significant biases: while all CorSym 95% confidence intervals (CIs) include zero, Pearson CIs in the original order exclude zero for both ASW and MXL. Thus, the observed order biases in ANCESTOR outputs are large enough to result in large and significant biases in ancestry correlation estimates.

**Table 2.** Two-way ANCESTOR biased ordering and its impact on Pearson correlation bias.

| Population | Biased order proportion |  |  | Correlation estimates |  |  |  |
| --- | --- | --- | --- | --- | --- | --- | --- |
| | $P1 \geq P2$ | $\hat{f}$ | P-value | $r_c$ | $\bar{r}_p^{\text{rand}}$ | $r_p$ | $r_p - r_c$ |
| ACB | 16 | 0.8 | 0.012 | -0.197 | -0.218 | -0.02 | 0.177 |
| ASW | 13 | 1 | 2.44e-04* | -0.139 | -0.155 | 0.784 | 0.923 |
| CLM | 24 | 0.686 | 0.041 | -0.114 | -0.116 | -0.034 | 0.08 |
| MXL | 21 | 0.656 | 0.11 | -0.281 | -0.408 | -0.377 | -0.096 |
| PEL | 21 | 0.6 | 0.311 | 0.066 | 0.07 | 0.113 | 0.047 |
| PUR | 22 | 0.629 | 0.175 | 0.199 | 0.215 | 0.253 | 0.054 |
| Total | 117 | 0.688 | 1.02e-06* | - | - | - | - |

\*Test is significant at 0.05 level after Bonferroni correction for 7 tests ( $p < 0.00714$ ). $\bar{r}_p^{\text{rand}}$ is mean Pearson estimate over 100 randomizations. Correlations not calculated for total since they are subject to Simpson’s paradox, since different populations have different ancestry averages.

**Figure 3.**
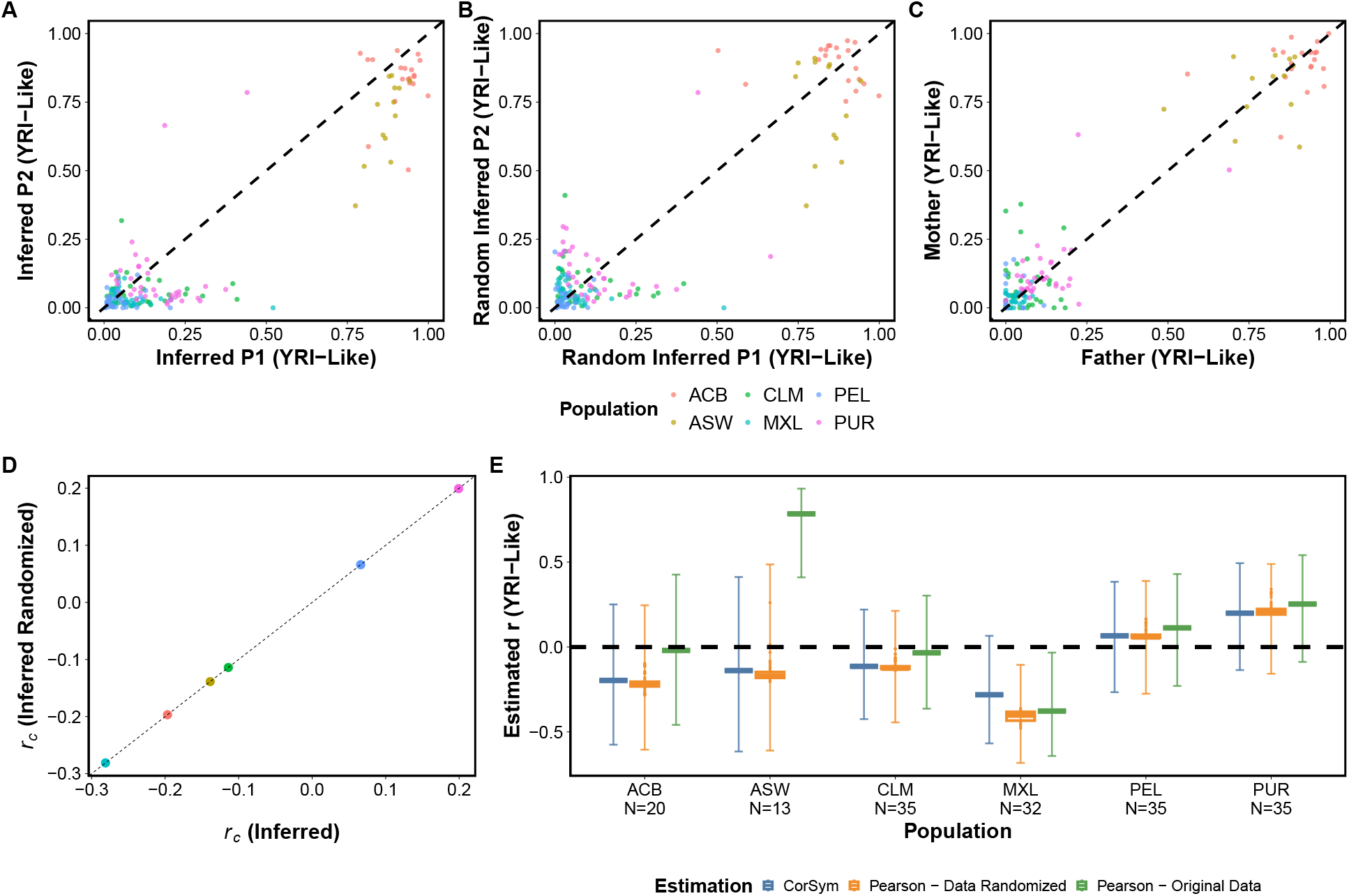
Biased ordering of inferred parental admixture biases assortative mating correlation estimates. **A-C)** Comparing YRI-like ancestry between parents from different methods. **(A)** ANCESTOR ordered as given. Note P1 tends to have larger values than P2 (more points below diagonal), which later causes a correlation estimation bias. **(B)** ANCESTOR after randomizing pairs. **(C)** ADMIXTURE (true ancestry) ordered by sex. **D)** CorSym calculates the same correlation for both the ANCESTOR given and randomized orders. **E)** CorSym and Pearson generally agree when order is randomized, while Pearson estimates are highly biased when using the order given by ANCESTOR.

We also performed three-way inference with ANCESTOR, which resulted in biased ordering but only for one ancestry. Averaging parent estimates (to avoid having to align parents for the moment), Non-YRI-like ancestry from two-way analysis was split here between IBS-like and NAM-like ancestries, except in ACB and ASW this component is primarily IBS-like, as expected (Fig. S5). The pooled order proportions were significantly biased only for NAM-like ancestry (Table 3, Fig. S6A,C,E). Of note, ancestry references were ordered as NAM, IBS, and YRI in the input to ANCESTOR. We also tested the altered reference order of IBS, NAM, YRI, and no significant order biases were observed (Table S3). Thus, ANCESTOR’s order biases in three-way analysis depend on reference order.

**Table 3.**
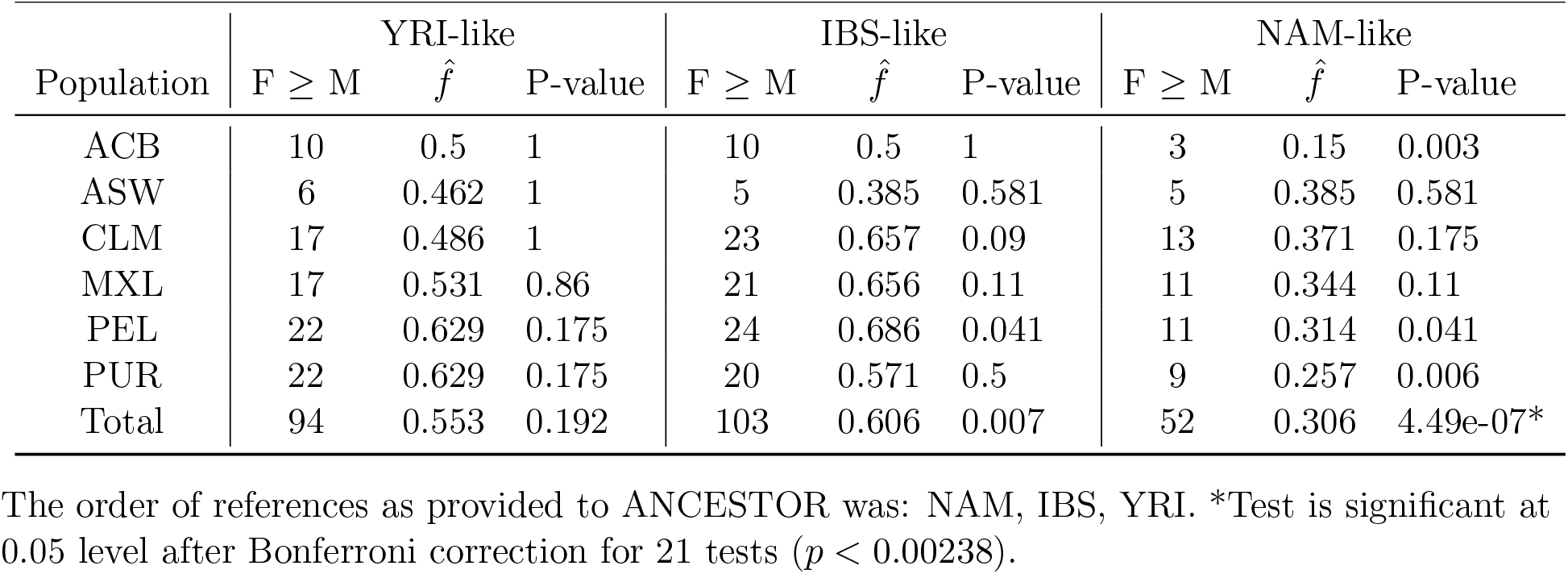
Three-way ANCESTOR biased order proportions.

#### 3.2.3 ANCESTOR experiences an artifact specific to three-way analysis

Our three-way analysis also had large errors in some individuals, which were not observed in the two-way analyses. An artifact in ANCESTOR caused some children to have both inferred parents arbitrarily placed near 50% ancestry for the first two references in the input (IBS-like and NAM-like), while placing the third component (YRI-like) at zero (Figs. S5 and S6A,C,E). This artifact depends on the order of the reference populations, since it shifts to IBS-like and YRI-like ancestries when they are placed first (Fig. S7). We see broad agreement between estimates of child global ancestry by RFMix (the step before ANCESTOR) and ADMIXTURE (Fig. S8A,D,G) and between ADMIXTURE mean parent and child estimates (Fig. S8B,E,H), with no similar artifact present. In contrast, a portion of children disagree between their true ancestries and their ANCESTOR mean parent estimates precisely as the artifact manifests, by incorrectly having ancestry near 0.5 for the first two ancestries and 0 for the third one (Fig. S8C,F,I). Thus, ANCESTOR is the source of these large errors present in a subset of individuals.

#### 3.2.4 ANCESTOR overestimates divergence between parent ancestries

To examine overall ANCESTOR accuracy, inferred and true parental ancestries were compared. We observed higher mean differences between parent ancestries compared to their true ancestry in two-way analysis (Fig. 4A) and three-way analysis except for YRI-like ancestry, for both input reference orders tested (Figs. S6 and S7). Since average parent inferred and true ancestry broadly agree in 2-way (Fig. 4B) and 3-way analysis (except for a few parents suffering from the artifact; Fig. S8), this error is restricted to overestimating the difference between parent ancestries. For that reason, we call it the divergence bias, which by itself is expected to reduce parent correlation estimates toward zero.

**Figure 4.**
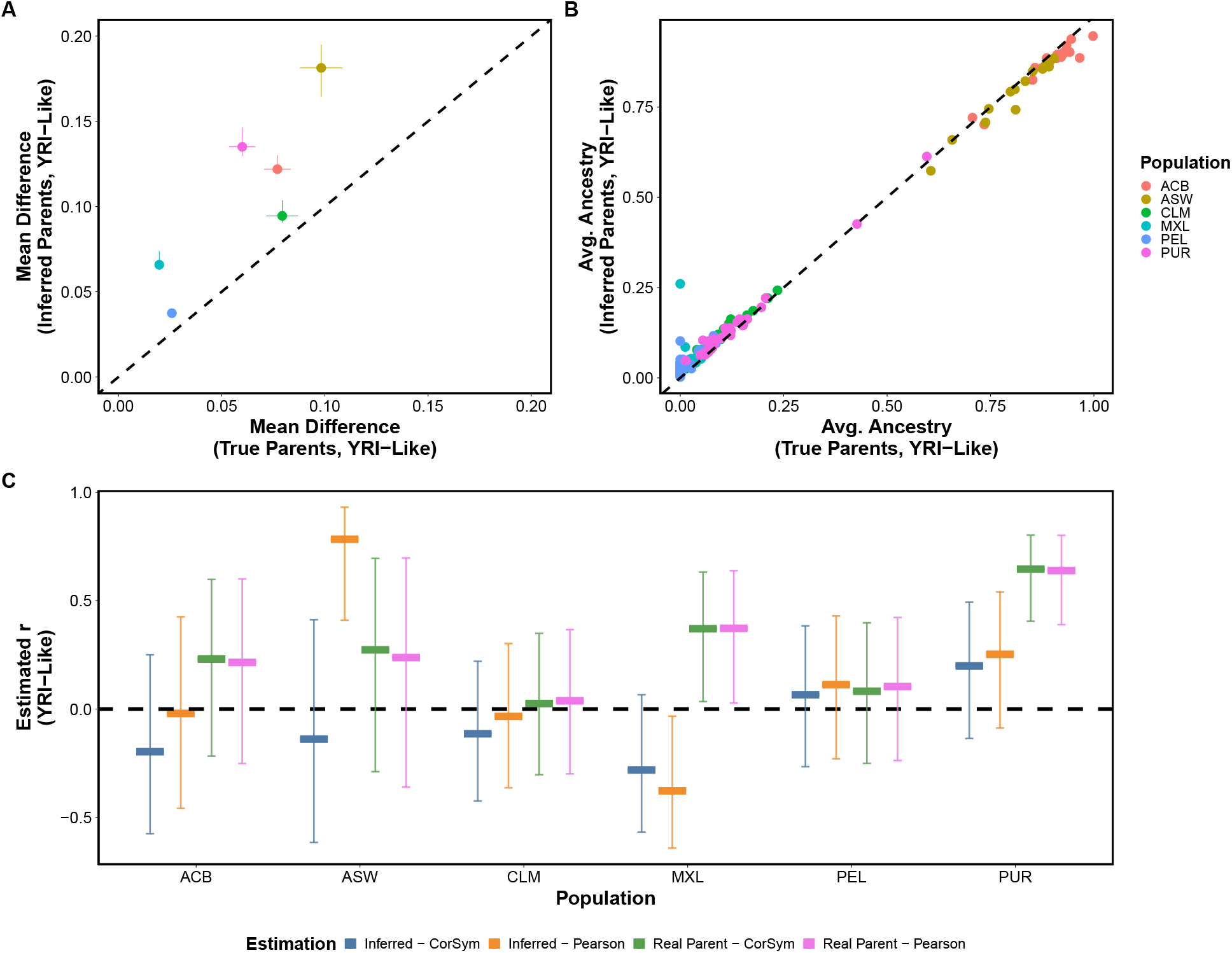
Ancestry correlations between true and inferred parents differ. Two-way ANCESTOR (inferred) and ADMIXTURE (true) ancestry is used here. **A)** Inferred parents show consistently higher ancestry differences than true parents. **B)** Inferred and true parental ancestry averages are similar. **C)** Inferred (same as Fig. 3E) and true parental ancestry correlations differ more than comparisons between correlation estimators.

Lastly, we look at the correlation estimates obtained from these data. In the 2-way analysis, comparing estimates for each combination of the two correlation estimators and between true sex-ordered and ANCESTOR-inferred ancestries, true ancestry estimates are most concordant between estimators (Fig. 4C, Table S4), agreeing with sex-ordered data being exchangeable. In particular, although the ANCESTOR order bias generally inflates Pearson estimates compared to CorSym, the divergence bias between parents resulted in a stronger bias toward zero: among all ANCESTOR-derived estimates, only ASW and PEL Pearson estimates were larger than true ancestry correlations. Focusing on CorSym only, while all ANCESTOR CIs include zero, true ancestry estimates are significantly positive in MXL and PUR.

In three-way analysis we also see agreement between estimates applied to real ancestries, and evidence of order biases in NAM-like ancestry, but in this data we saw more cases of ANCESTOR-derived estimates exceeding real ancestry correlations (Fig. 5 and Table S5). Focusing on significantly non-zero CorSym estimates, ANCESTOR gave overestimated correlations compared to real ancestry in YRI-like ancestry for ASW and PEL, in IBS-like for PUR, and in NAM-like for ACB, ASW and PUR. In contrast, CorSym applied to real ancestries identified significantly non-zero estimates that ANCESTOR did not in YRI-like ancestry for MXL and PUR (same as 2-way results), in IBS-like for MXL and PEL, and in NAM-like for MXL and PEL. Lastly, both CorSym estimates were significantly non-zero for PUR in IBS-like and NAM-like, though the estimates are quite different between ANCESTOR and true ancestry. The real ancestry results point to strong evidence of assortative mating in XML, PEL, and PUR. Significance was not reached in ACB, ASW, and CLM, but note the first two also had considerably smaller sample sizes. Thus, ANCESTOR estimates exhibit multiple biases with opposite directions that together result in unreliable parental ancestry correlation estimates.

**Figure 5.**
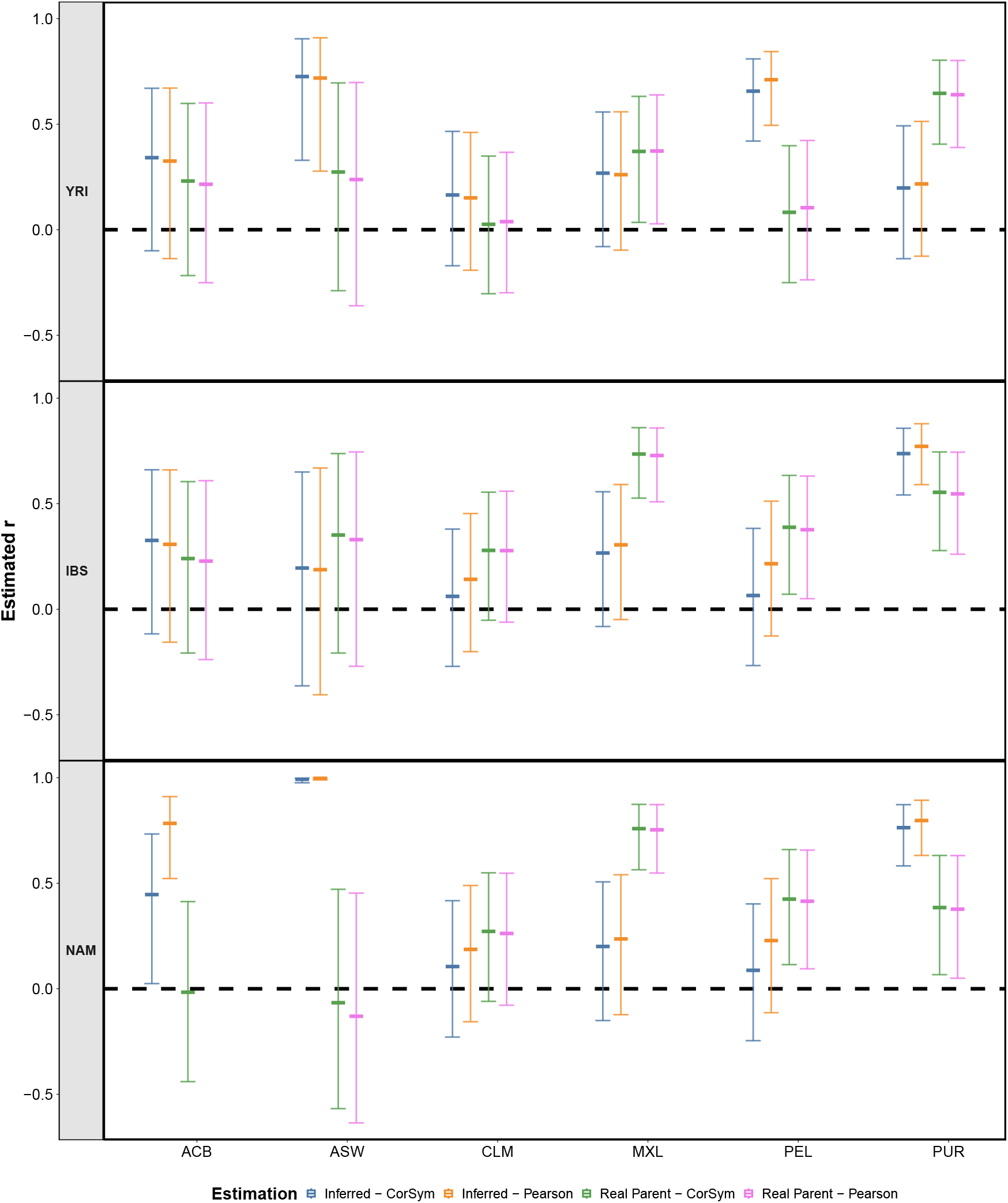
Three-way true and ANCESTOR-inferred correlations disagree substantially. Compared to two-way, three-way ANCESTOR outputs have a diminished order bias, but comparable and substantial divergence bias, as well as new artifact errors, all causing correlation biases. Pearson had NA correlation for NAM-like ACB in real parents because all parents of one sex had none of this ancestry (zero variance).

## 4 Discussion

In this work we considered a pair of exchangeable variables, which by definition are not meaningfully ordered, and which arise in the study of assortative mating and other fields. The specific order of the data incorrectly alters standard correlation estimates. Ordered data frequently induce an upward bias in standard correlation estimates that could lead to misleading results, as in the assortative mating example where the correlation estimate is used to make inferences on the mating structure of a population. To fix this bias, we introduced CorSym, a correlation estimator that assumes identical marginal distributions, and by design does not depend on the order within pairs. We considered how order bias impacts Pearson estimates and compared it with CorSym through theory and simulated data. Lastly, we present the real example where biased order manifests and motivated this work, namely parent ancestry inference with ANCESTOR, and demonstrated Pearson estimates are greatly affected by ordering.

Most correlation analyses involve two completely asymmetrical variables, such as temperature and the number of cars on the road, where it makes no sense to consider reordering the two variables in any one pair only. However, there are questions involving the correlation of exchangeable paired data, where order is arbitrary and uninformative, but as we showed here, standard estimators are confounded by biased pair orders. An example of such exchangeable data is paired measurements of the same variable, where there is no additional information that may define an informative order for these measurements. The application we focused on is parental ancestry, where no ordering information is available such as sex. Other examples could be temperature readings between random locations paired by proximity, or the amount of sleep couples receive measured across thousands of pairs. In these cases, the correlation of the paired information is important, but not the identity and order of the two variables.

Our analysis of admixed 1000 Genomes trios points to fathers and mothers having similar ancestry distributions in a given population, so they can be treated as exchangeable. However, Hispanic and African-American populations are known to have experienced sex-biased admixture in the past [29, 30, 31, 32, 8]. In the initial generations, when large numbers of immigrants arrive and mate with a local population with sex-biased patterns, parents do not have exchangeable ancestry values. Nevertheless, child ancestries at autosomes will be averages of their parents, while their sex is assigned randomly, so in the following generation men and women have exchangeable ancestry proportions, absent further sex-biased migration. Non-autosomes, not considered in this work, do have ancestry differences under sex-biased admixture that allows it to be detected. Thus, our results are consistent with present day admixed populations being at a steady state where ongoing sex-biased admixture is negligible, justifying our assumption of exchangeability.

If the variables of interest are truly exchangeable, order biases can arise in many ways. Heuristics, mental shortcuts humans use to make faster decisions [33, 34, 35] and are often made without awareness [36], can result in order biases. For example, a hypothetical researcher entering data could use a heuristic where they always start with two digit sleep times over one digit, because they expect it to take slightly longer. An extreme order could also be imposed to misguidedly ensure that order is deterministic, not knowing that such a choice maximizes correlation biases. In addition to those hypothetical scenarios, we found that the ANCESTOR software outputs partially ordered parent ancestry pairs. The resulting biased correlation estimates are immediately concerning, affecting our inference of the presence and strength of assortative mating [37, 38, 39].

Beyond biased parent order, we also found that ANCESTOR inferred parents diverge in ancestry more than true parents, and some samples suffered from an inference artifact resulting in equal ancestry for two of the three ancestries. Order and divergence biases, and the artifact, all resulted in poor correlation accuracy overall, but each bias has an opposite direction that made the overall effect hard to predict. This work addressed order bias, but addressing the parents’ divergence bias and the artifact remain the topic of future work.

Overall, we identified an important scenario where Pearson correlation estimates are often upwardly biased, namely when order within each pair is arbitrary. The Pearson estimator assumes that order is meaningful, so it was not designed for this scenario in mind. Biased orders can arise in many scenarios, and could even be crafted maliciously to selectively exaggerate correlations as desired. One option to mitigate this problem is to randomize the variable order within each pair, but the result is no longer deterministic. To ensure unbiased and deterministic correlation estimates for exchangeable variables, we recommend using CorSym. On the other hand, applying CorSym to non-exchangeable data will result in its own biases. Therefore, pair exchangeability has to be known given the study design, established empirically as we did for real parent ancestries ordered by sex, or ideally both.

## Acknowledgments

Thanks to Dashiell Massey and Amy Goldberg for making us aware of the order bias in ANCESTOR outputs and its effect on Pearson correlation estimates, as well as other feedback on this work.

## Data and code availability

CorSym and utilities to bias and randomize pair orders, and to statistically test for order bias, are implemented in the R package `corsym`, available on GitHub at https://github.com/OchoaLab/corsym.

The data and analysis code generated during this study are available on GitHub at https://github.com/OchoaLab/corsym-project.

## Author contributions

GK: Methodology, Software, Validation, Investigation, Data Curation, Writing.

AO: Conceptualization, Methodology, Software, Formal Analysis, Writing, Supervision.

## Appendices A Derivation of covariance of extreme order variables

To proceed, recall the following properties of the mimimum *L* and the maximum *H* of *{X, Y}*: *L* + *H* = *X* + *Y* and *LH* = *XY*. Using the definition of covariance and *LH* = *XY*, we relate both covariances:

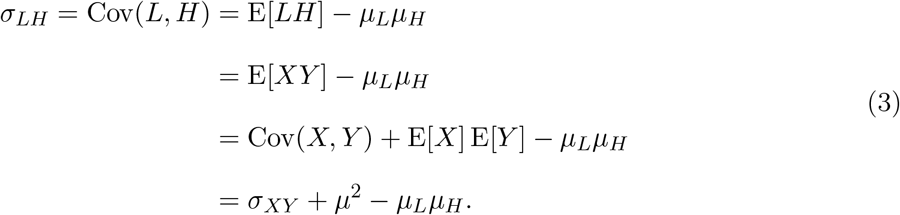

From *L* + *H* = *X* + *Y* it follows that

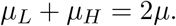

Solving for *µ*_*H*_ and plugging into Eq. (3), we get

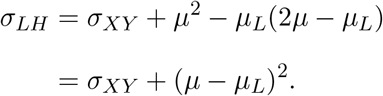

Therefore, since the square is always non-negative, the covariance of *L* and *H* is always greater or equal than that of *X* and *Y* : *σ*_*LH*_ ≥ *σ*_*XY*_.

As correlations, the previous equation becomes the first part of Eq. (2), reproduced here:

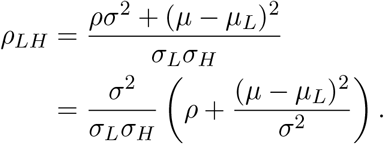

Expanding the variance of *L* + *H* = *X* + *Y*, we get that

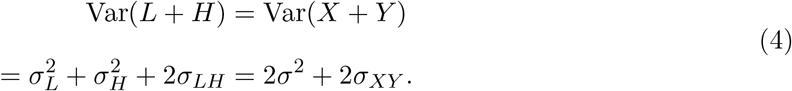

Since *σ*_*LH*_ ≥ *σ*_*XY*_, combining with Eq. (4) and applying the geometric-arithmetic mean inequality to the non-negative 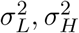, we get that

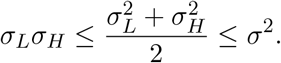

Therefore, since 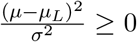 and 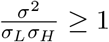, it follows from Eq. (2) that *ρ*_LH_ ≥ *ρ* when *ρ_LH_ >* 0, as desired, but the argument does not work for negative *ρ*_*LH*_ in general.

Writing the covariances in Eq. (4) as correlations, and solving for *ρ*_*LH*_, we get that

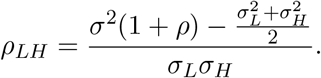

We wish to show that *ρ*_*LH*_ ≥ *ρ*, which is equivalent to showing that

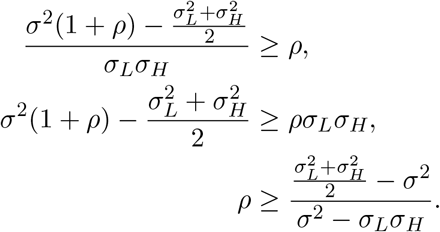

Note that to preserve the inequalities, we only multiplied and divided by positive terms (*σ*_*L*_*σ*_*H*_ and *σ*^2^ − *σ*_*L*_*σ*_*H*_), assuming no variances are zero and *σ*^2^ *> σ*_*L*_*σ*_*H*_ strictly (if *σ*^2^ = *σ*_*L*_*σ*_*H*_, then *ρ*_*LH*_ ≥ *ρ* follows directly from Eq. (2) as desired). From here it is clear that when *σ*_*L*_ = *σ*_*H*_, the inequality simplifies to *ρ* ≥ −1, which always holds, so *ρ*_*LH*_ ≥ *ρ* will also hold. However, if *σ*_*L*_≠ *σ*_*H*_, then applying the geometric-arithmetic inequality again, we see that

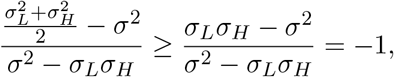

so unfortunately that term may now exceed *ρ* when *ρ* is very close to −1. Indeed, we found a counterexample where this occurs (Fig. S2).

## B Expectation calculations for baseline estimators

First we will need that

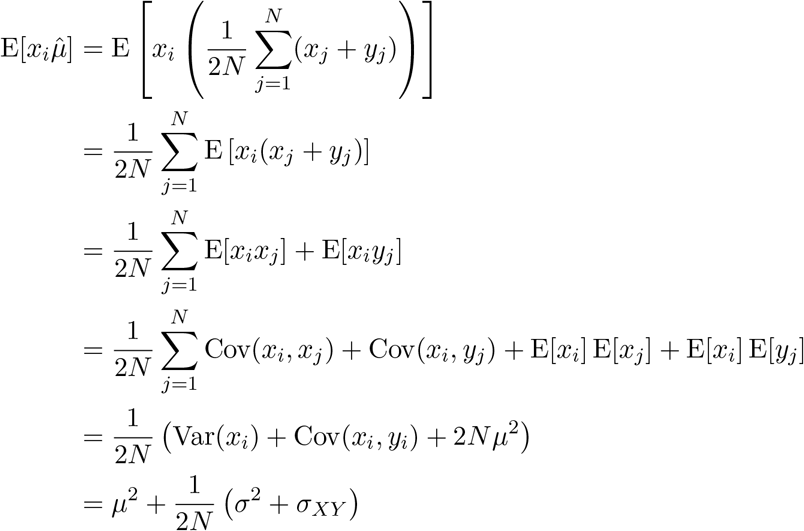

where we assumed that the covariance of different pairs (*j* ≠*i*) is zero. Therefore, 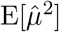 has the same value:

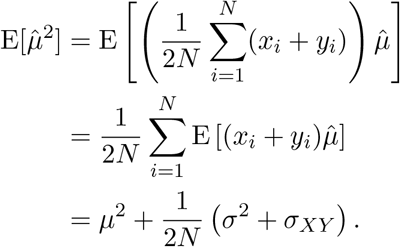

Next, we obtain that

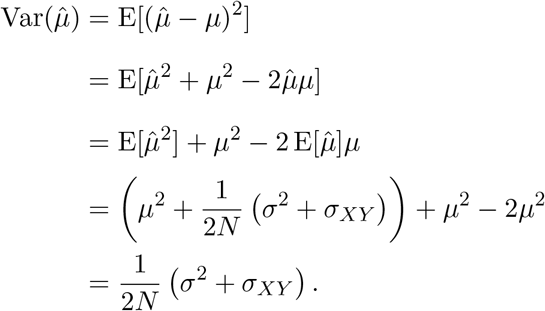

It follows that

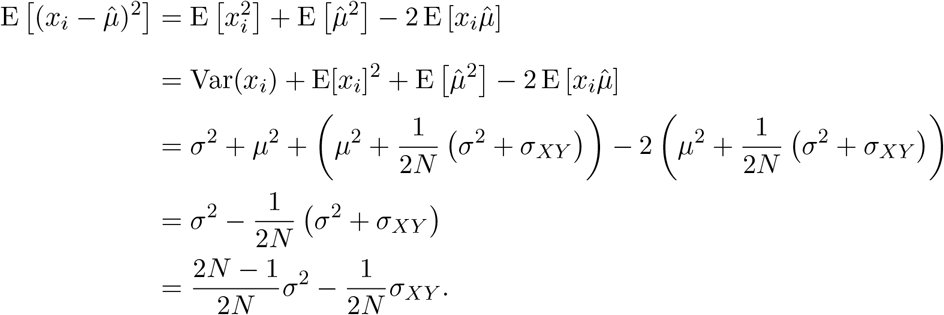

Lastly, we obtain the expectation of the base sample variance,

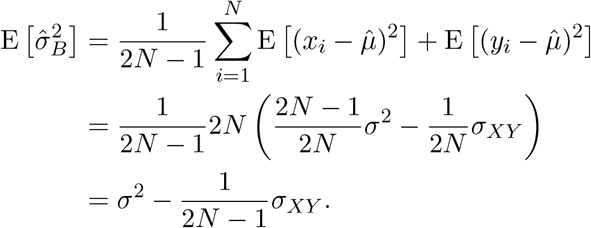

For the base sample covariance, we have

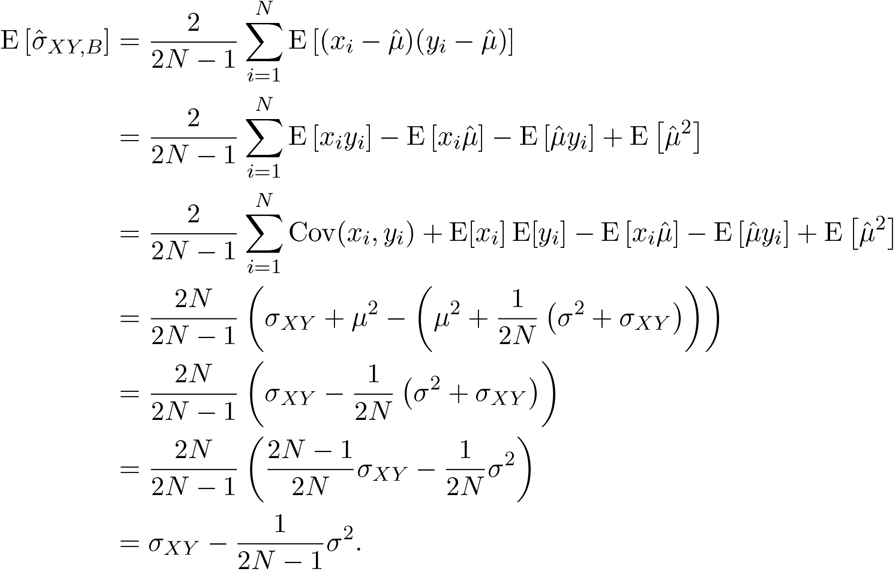

**C Solving system of equations to unbias base estimators**

The system of equations we desire to solve is given using matrix algebra by

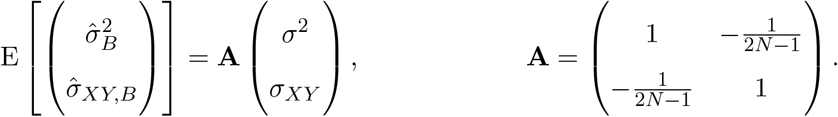

The inverse of the above matrix is

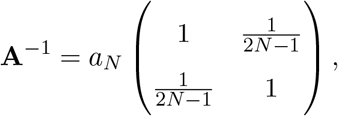

where *a*_*N*_ is the inverse of the determinant and matches what is given in the main text:

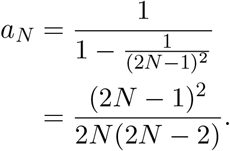

The unbiased estimators are given by

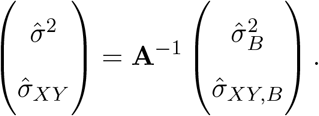

It follows that these estimators are actually unbiased:

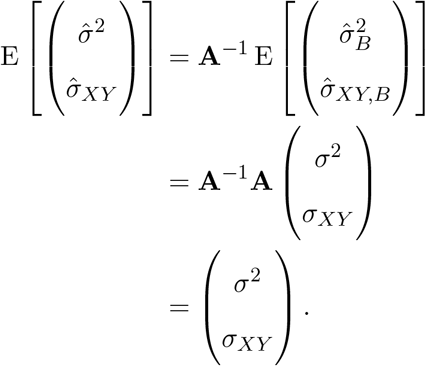

## S1 Supplementary tables and figures

**Table S1.**
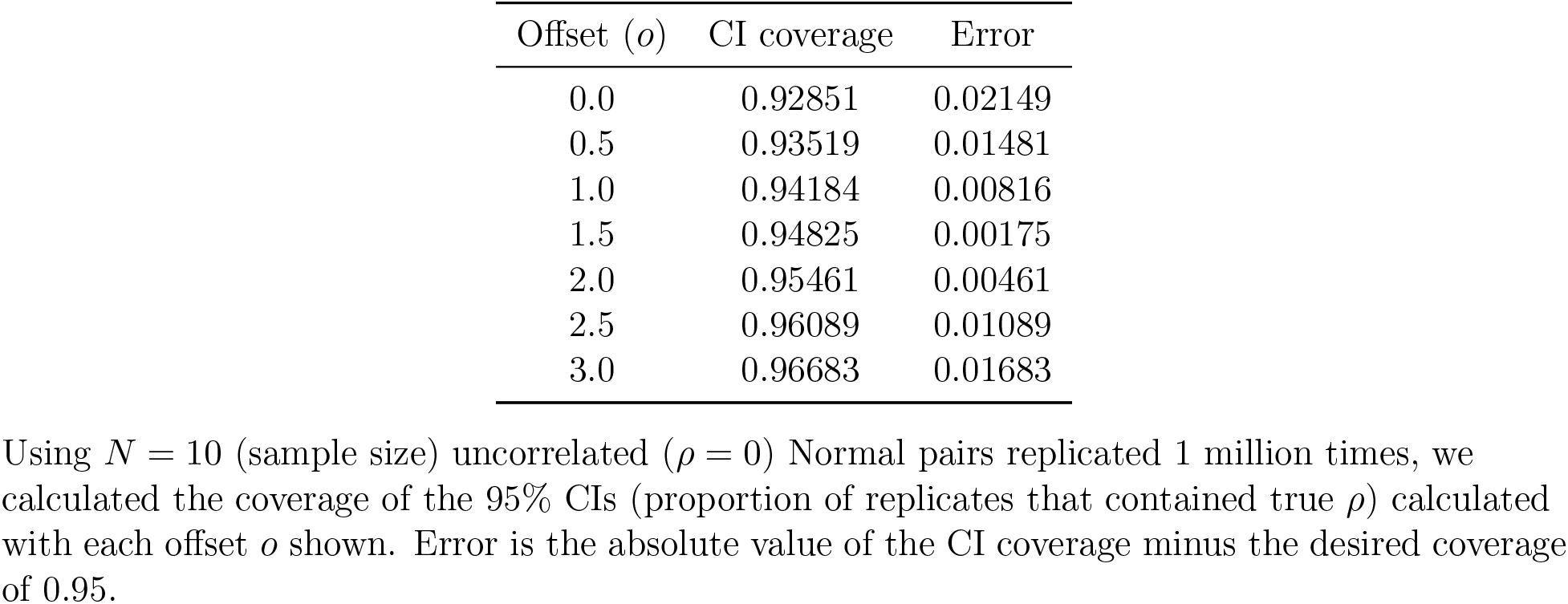
Fitting CorSym CI offset using simulations.

**Table S2.** Kolmogorov-Smirnov two-sided p-values testing if father and mother true ancestry distributions are different.

| Population | YRI-like | IBS-like | NAM-like |
| --- | --- | --- | --- |
| ACB | 0.336 | 0.336 | 0.487 |
| ASW | 0.588 | 0.588 | 0.096 |
| CLM | 0.836 | 0.690 | 0.874 |
| MXL | 0.418 | 0.830 | 0.828 |
| PEL | 0.621 | 0.977 | 0.977 |
| PUR | 0.979 | 0.979 | 0.979 |

No test was significant at the 0.05 level after Bonferroni correction for 18 tests ( $p < 0.0028$ ).

**Table S3.** Three-way ANCESTOR biased order proportions under alternate reference order.

| Population | YRI-like |  |  | IBS-like |  |  | NAM-like |  |  |
| --- | --- | --- | --- | --- | --- | --- | --- | --- | --- |
| | $P1 \geq P2$ | $\hat{f}$ | P-value | $P1 \geq P2$ | $\hat{f}$ | P-value | $P1 \geq P2$ | $\hat{f}$ | P-value |
| ACB | 16 | 0.800 | 0.012 | 4 | 0.200 | 0.012 | 11 | 0.550 | 0.824 |
| ASW | 9 | 0.692 | 0.267 | 4 | 0.308 | 0.267 | 4 | 0.308 | 0.267 |
| CLM | 10 | 0.286 | 0.017 | 26 | 0.743 | 0.006 | 13 | 0.371 | 0.175 |
| MXL | 12 | 0.375 | 0.215 | 12 | 0.375 | 0.215 | 21 | 0.656 | 0.110 |
| PEL | 16 | 0.457 | 0.736 | 13 | 0.371 | 0.175 | 22 | 0.629 | 0.175 |
| PUR | 18 | 0.514 | 1.000 | 19 | 0.543 | 0.736 | 16 | 0.457 | 0.736 |
| Total | 81 | 0.476 | 0.591 | 78 | 0.459 | 0.319 | 87 | 0.512 | 0.818 |

The order of references as provided to ANCESTOR was: IBS, YRI, NAM. No test was significant at 0.05 level after Bonferroni correction for 21 tests ( $p < 0.00238$ ).

**Table S4.** True Parent (ADMIXTURE) correlation estimates largely agree.

| Population | YRI-like |  |  | IBS-like |  |  | NAM-like |  |  |
| --- | --- | --- | --- | --- | --- | --- | --- | --- | --- |
| | $r_c$ | $r_p$ | $r_p - r_c$ | $r_c$ | $r_p$ | $r_p - r_c$ | $r_c$ | $r_p$ | $r_p - r_c$ |
| ACB | 0.231 | 0.215 | -0.015 | 0.24 | 0.228 | -0.012 | -0.016 | NA | NA |
| ASW | 0.273 | 0.238 | -0.036 | 0.352 | 0.33 | -0.022 | -0.066 | -0.13 | -0.064 |
| CLM | 0.026 | 0.038 | 0.013 | 0.279 | 0.278 | -0.001 | 0.272 | 0.262 | -0.01 |
| MXL | 0.371 | 0.373 | 0.002 | 0.735 | 0.728 | -0.007 | 0.759 | 0.753 | -0.006 |
| PEL | 0.082 | 0.104 | 0.022 | 0.388 | 0.377 | -0.011 | 0.425 | 0.415 | -0.01 |
| PUR | 0.646 | 0.64 | -0.006 | 0.554 | 0.546 | -0.007 | 0.385 | 0.377 | -0.008 |

Pearson NA results from one sex having all zeroes for the given ancestry, but the other sex had variation so CorSym estimates a non-NA value.

**Table S5.**
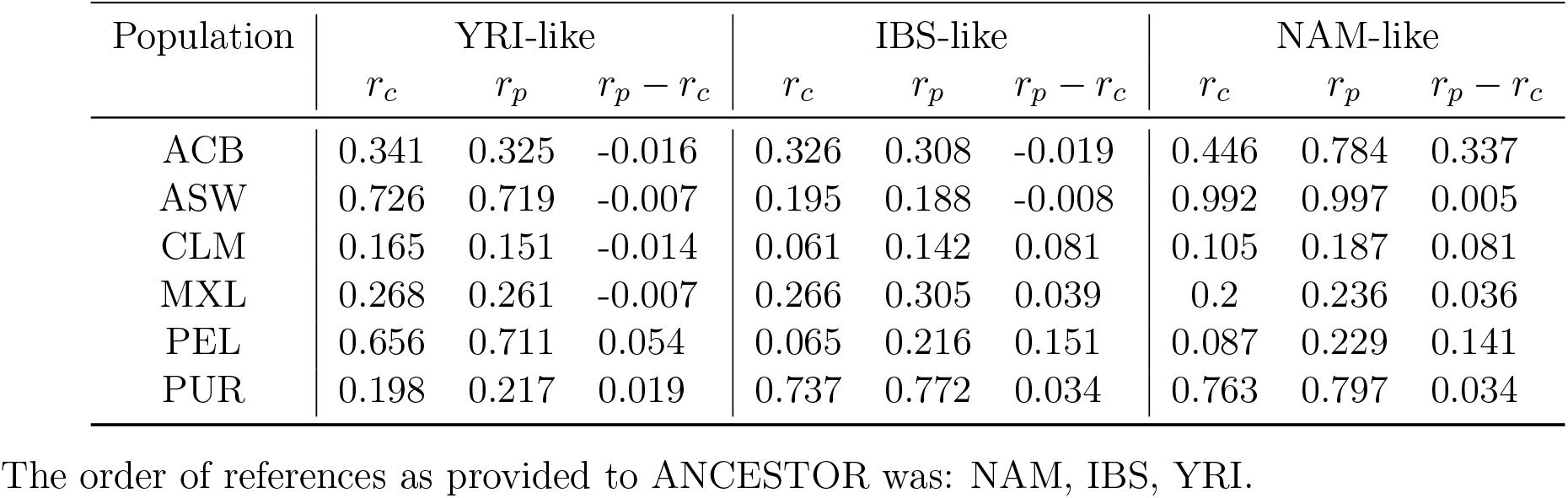
Three-way inferred (ANCESTOR) correlation estimates.

**Figure S1.**
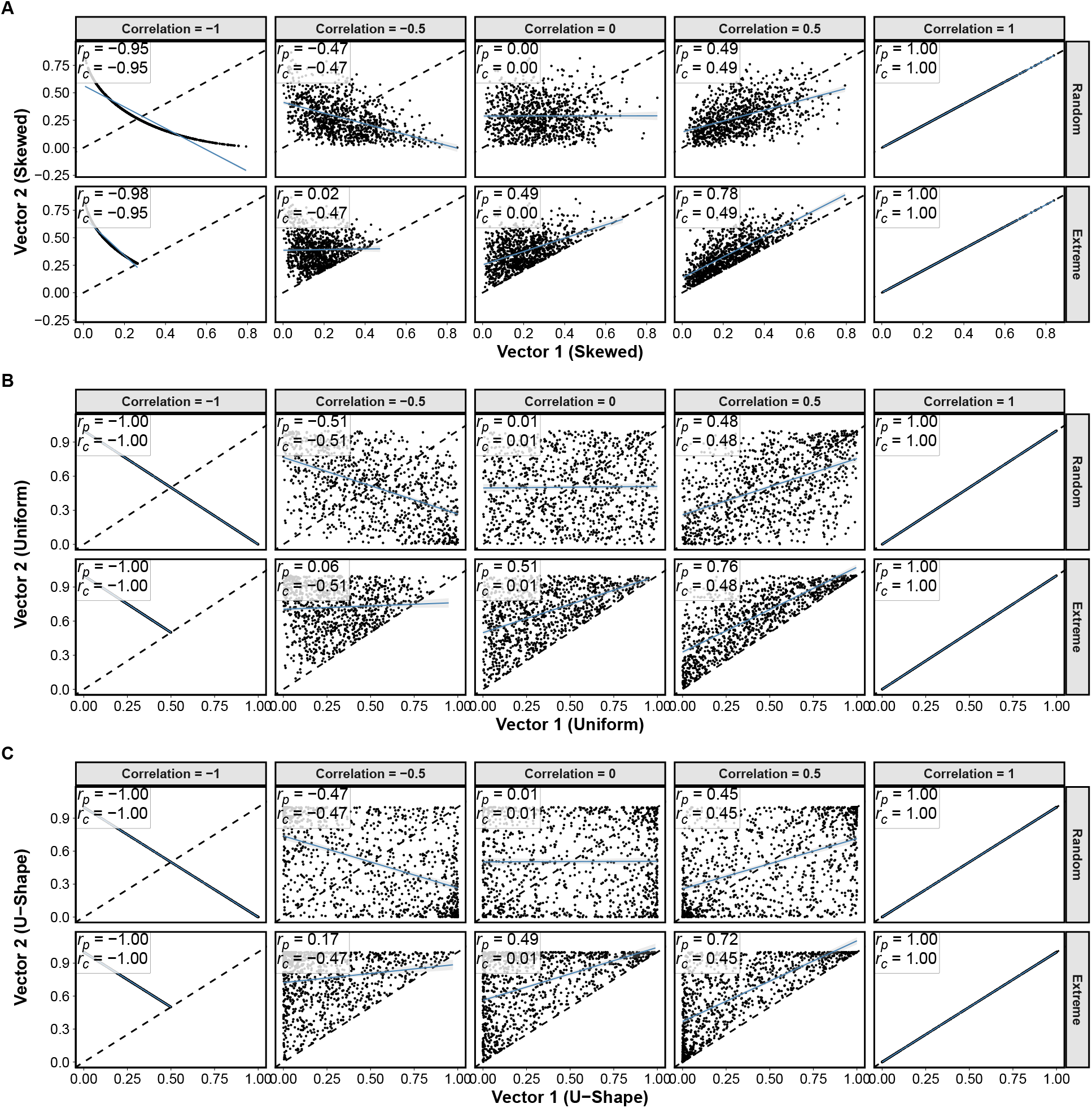
Order in non-normal simulated data affects Pearson but not CorSym estimates. **A)** Skewed, **B)** Uniform, and **C)** U-shaped multivariate distributions of exchangeable variables all show the same effect of ordering bias on Pearson’s estimates. Here the true correlation values (column titles) are those of the copula, which correctly specify the final correlation of the uniform data, and happens to work for U-shape as well, but is misspecified for Skewed. In those cases, random order estimates are unbiased for the true final correlation, appropriate to compare against the extreme order estimates.

**Figure S2.**
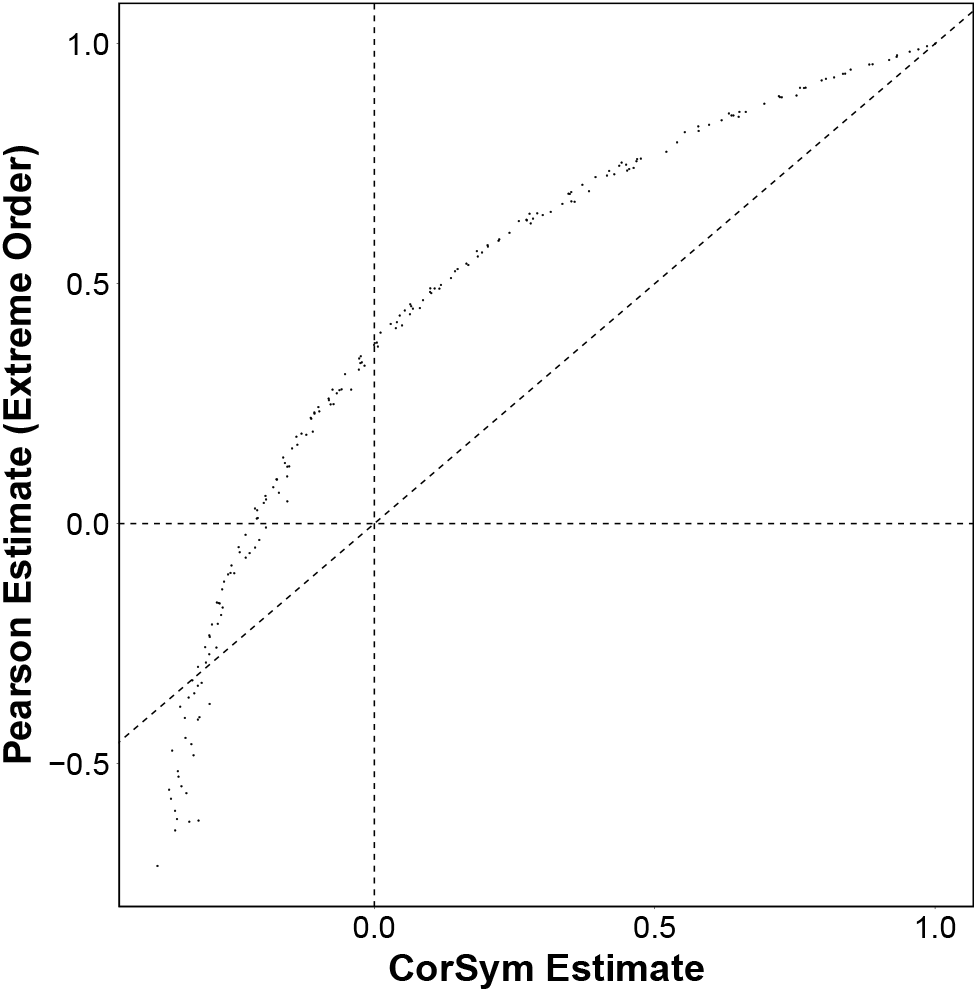
Highly skewed marginal distributions can result in downward Pearson biases under extreme order. We simulated bivariate Log Normal data, which is highly skewed, with *N* = 10, 000 pairs and correlation parameters between −1 and 1 in increments of 0.01, and for each case we ordered data extremely and plot the CorSym and Pearson estimates. Correlation parameters are for Normal data before exponentiation, so they do not equal the final correlations that CorSym estimates without bias. Note *ρ*_*LH*_ ≤ *ρ* occurs where the curve is below *y* = *x* (dashed diagonal line), which is a small portion at the lower left corresponding to highly negative correlations.

**Figure S3.**
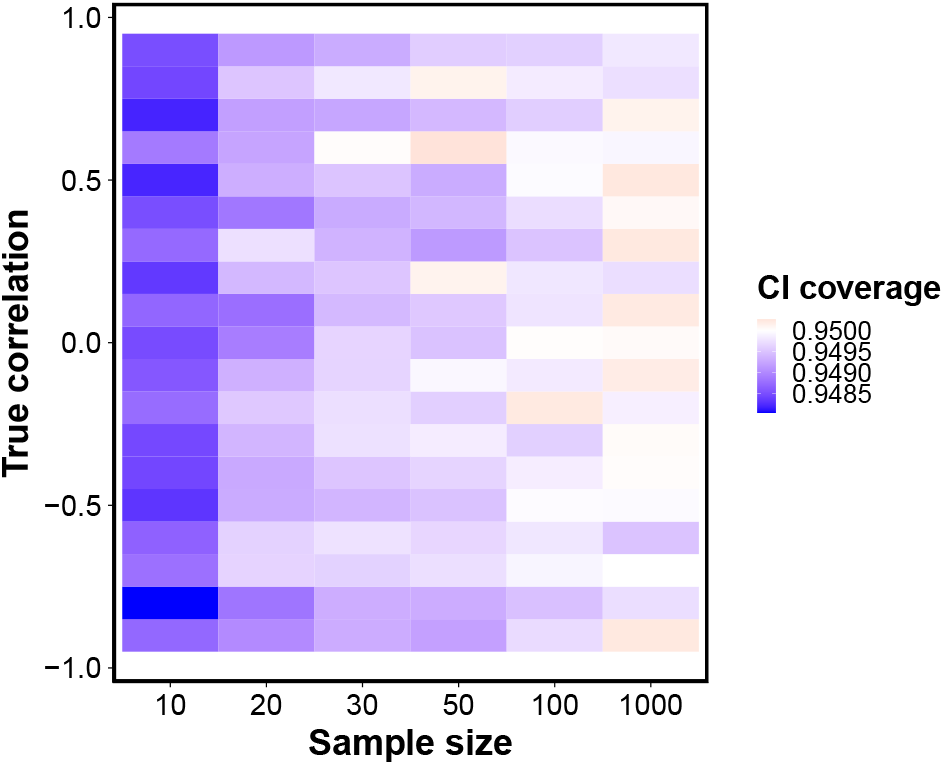
CorSym 95% confidence intervals (CIs) have the desired coverage in simulations. Bivariate Normal pairs with the sample sizes *N* and true correlations *ρ* shown in the x and y axes were used to estimate CIs 1 million times in each cell, and color shows the CI coverage, which is the proportion of times CIs contained the true parameter. Coverage error does not depend on *ρ*. Smaller sample sizes result in lower CI coverages than desired, but maximum errors are less than 0.002.

**Figure S4.**
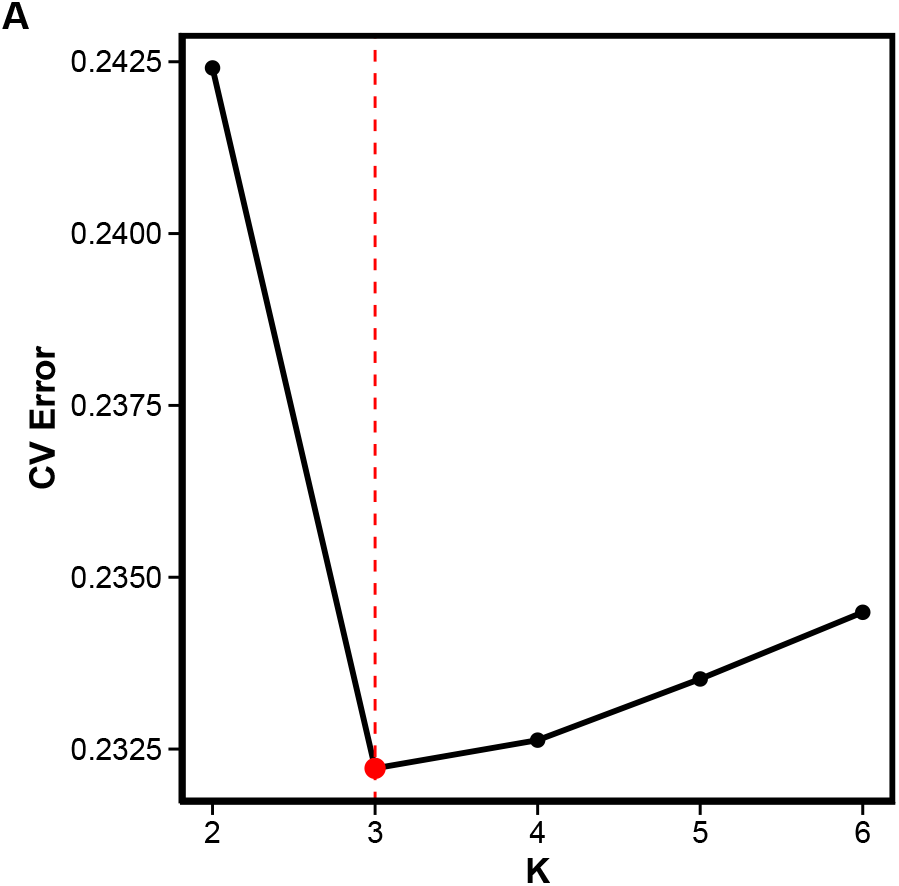
ADMIXTURE selection of number of ancestries using cross-validation. *K* = 3 is the best fitting admixture model in our merged dataset of admixed trios and references from 1000 Genomes and HGDP.

**Figure S5.**
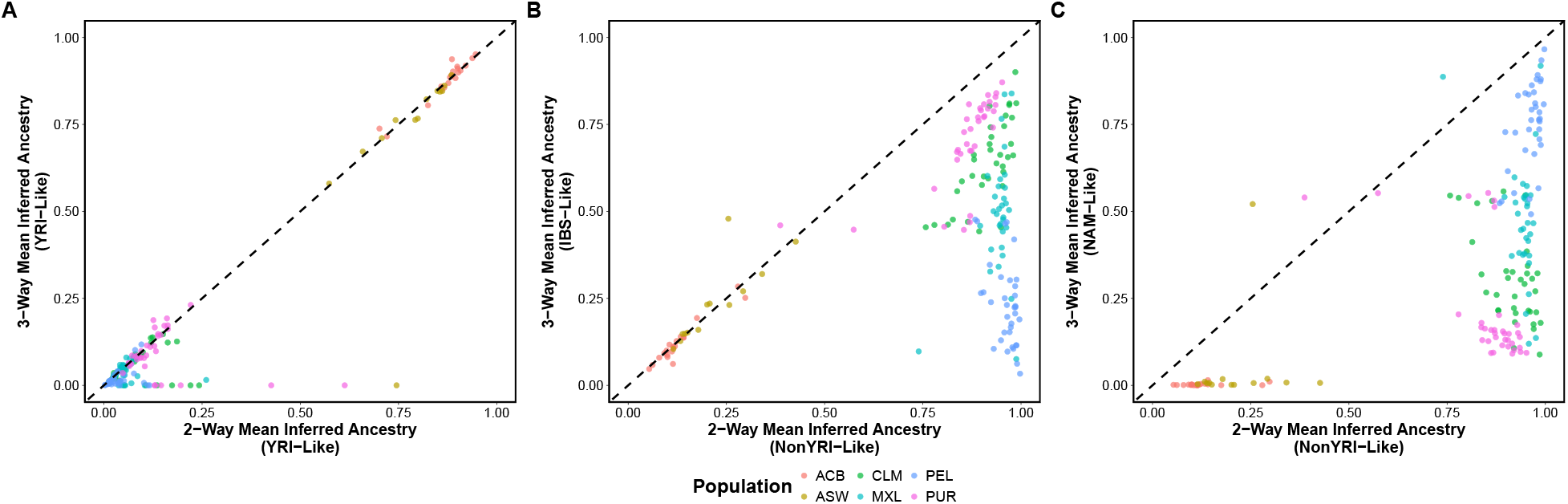
Comparison of parent-averaged ancestry estimates between two-way and three-way ANCESTOR outputs. **A)** The YRI component of two-way and three-way inference is generally equivalent, except for samples suffering the three-way artifact. **B**) The IBS component corresponds to the Non-YRI component for ACB and ASW populations. **C**) The NAM component correspond to the remainder of the non-YRI-like components in the Hispanic populations.

**Figure S6.**
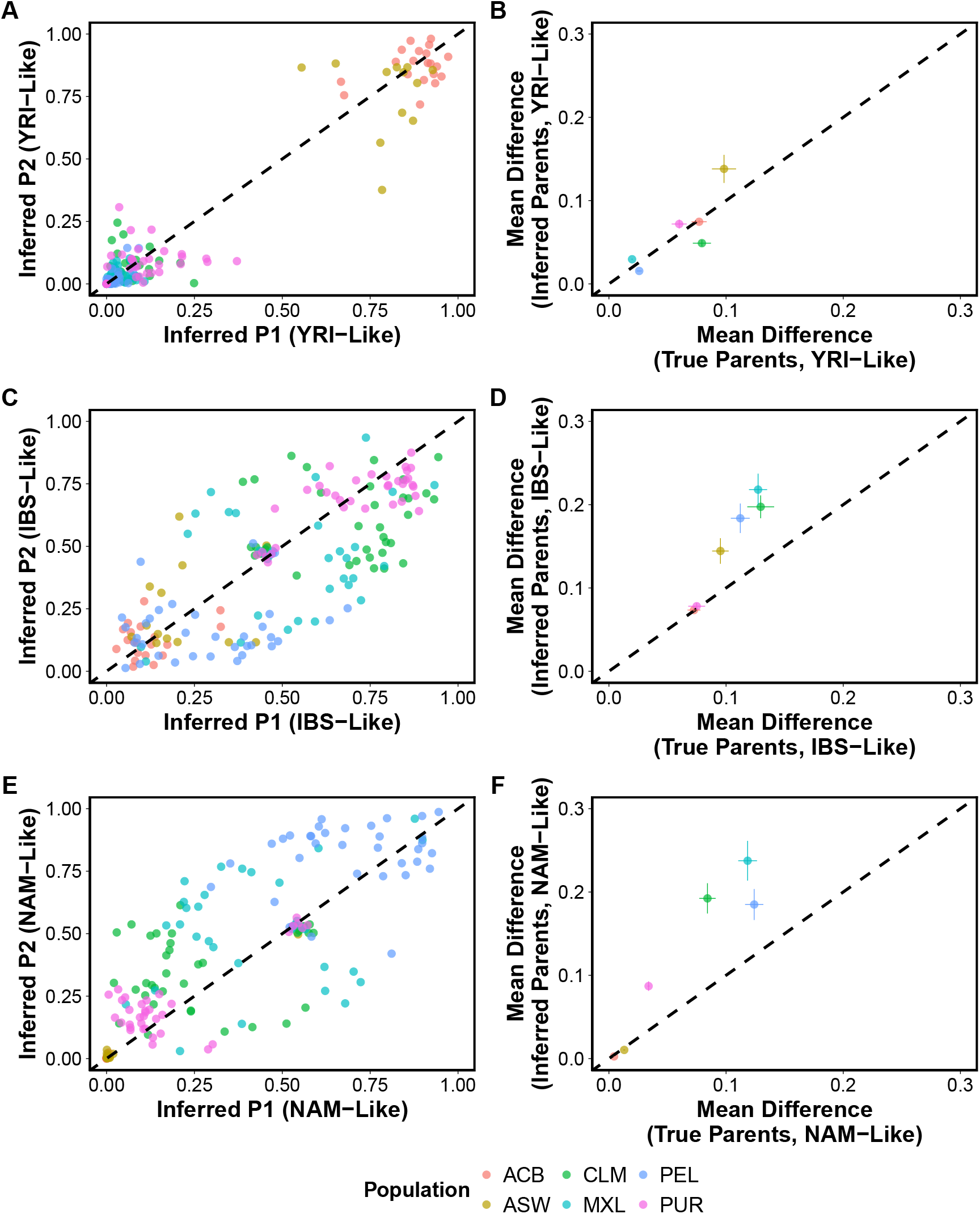
ANCESTOR three-way parent ancestry estimates differ more than those of true parents. **A,C,E)** Three-way analysis does not have as strong an order bias as two-way, but instead shows greater divergence (concentrations away from the diagonal), and also an artifact in IBS and NAM ancestries that results in many estimates near the middle point (0.5,0.5). **B,D,F)** Across all ancestries, inferred parents are more ancestrally different than real parents.

**Figure S7.**
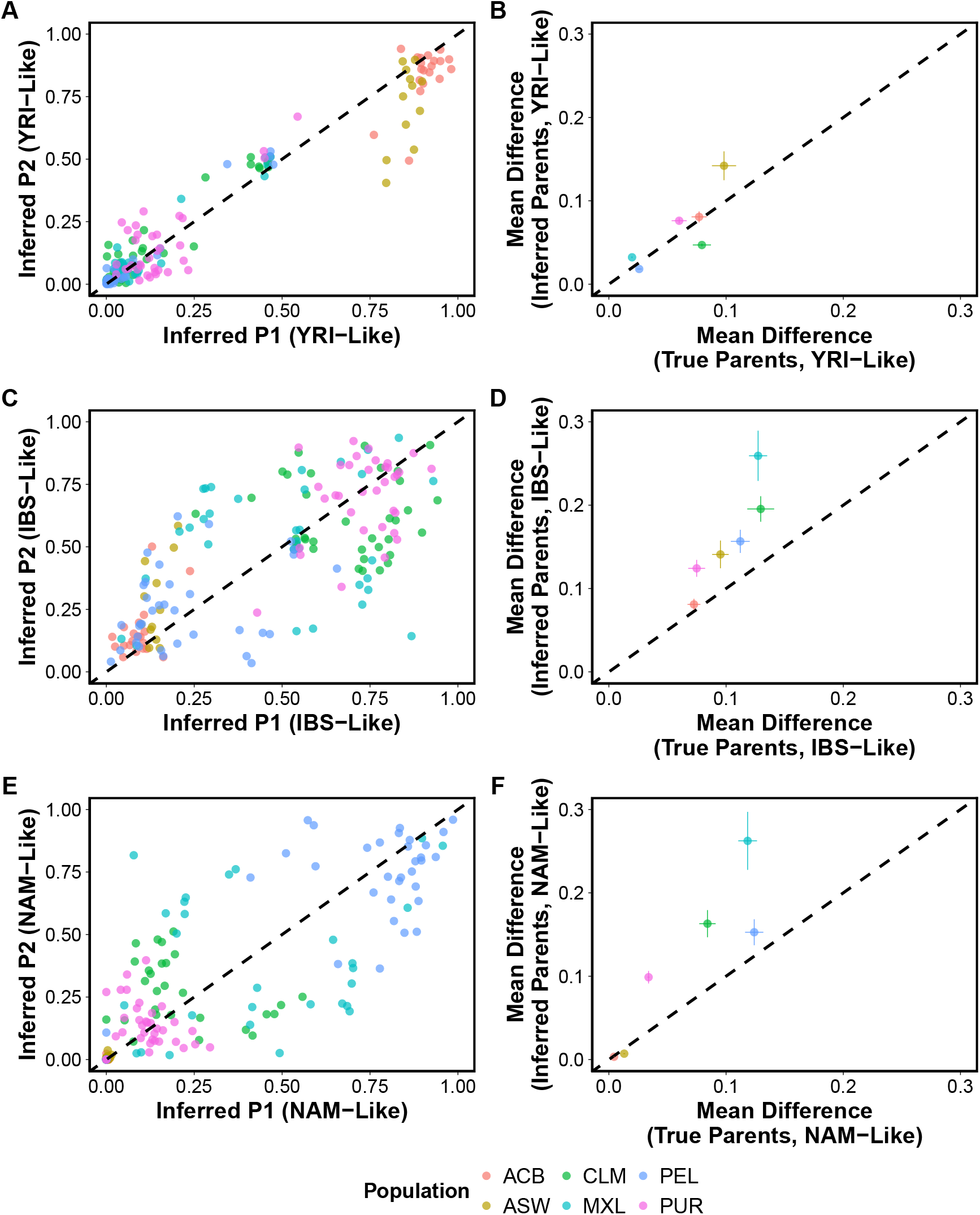
Order provided to ANCESTOR impacts artifact and order bias in three way analysis. The order of ancestries as provided to ANCESTOR was: IBS, YRI, NAM. This is a rerun of Fig. S6, where order of provided references changed. **A,C,E)** The reordered analysis shows similar divergence (concentrations away from the diagonal) to original analysis, and the artifact which was in IBS and NAM is now in IBS and YRI, suggesting that it is specified by the first two reference ancestries provided in the input. **B,D,F)** Inferred parents are more ancestrally different than real parents, similarly to previous results.

**Figure S8.**
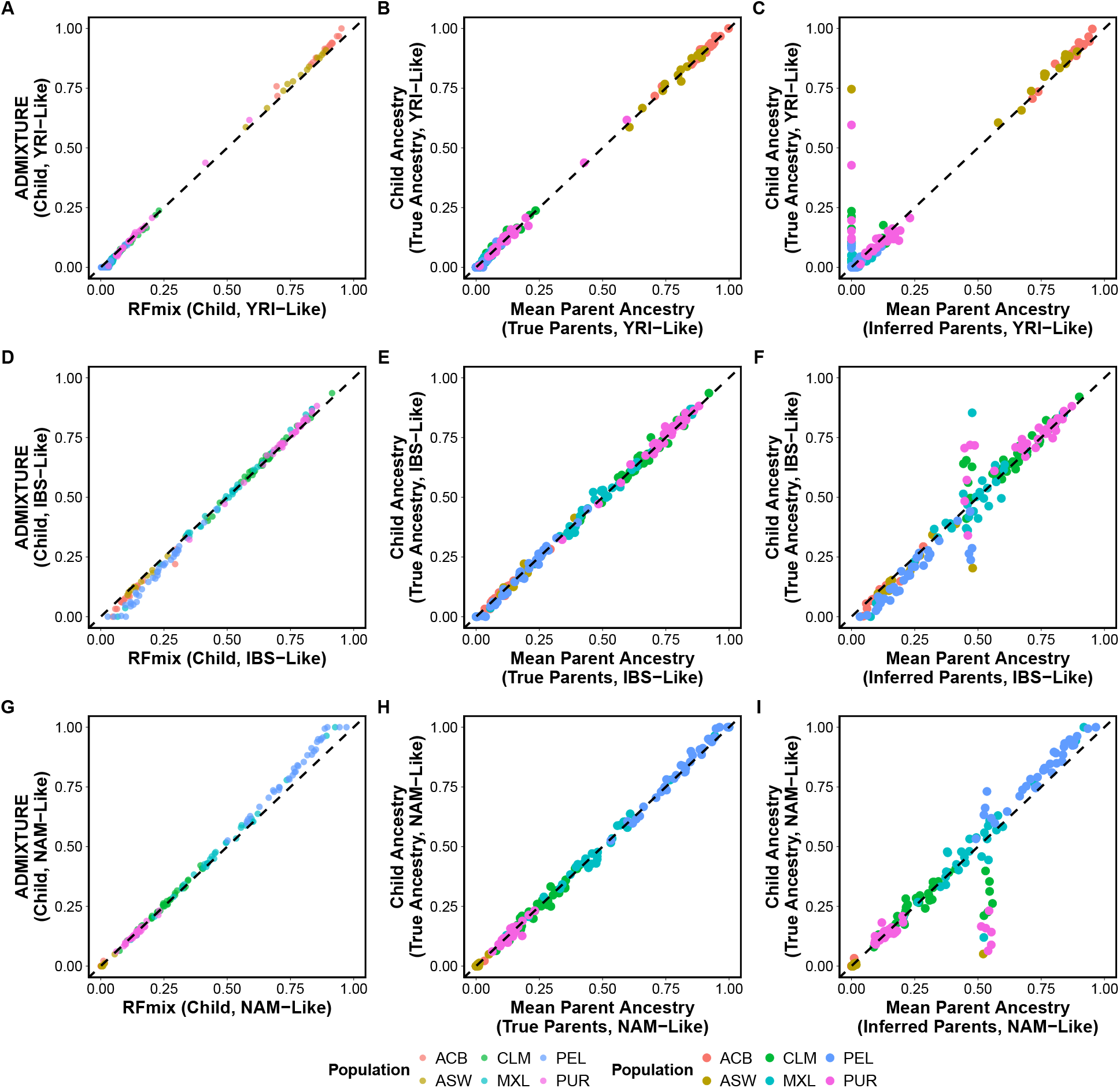
Three-way true and inferred parent ancestry means are similar to child’s ancestry. **A,D,G)** RFmix and ADMIXTURE child estimates largely agree, except for slight reductions for high NAM-like ancestry (made up by increases in low IBS-like) in RFmix. **B,E,H)** True mean parent ancestries closely follow children’s ancestry. **C,F,I)** ANCESTOR-inferred parents similarly agree on average with the input child data, except for data impacted by the reported artifact.

